# Autism-associated CHD8 keeps proliferation of human neural progenitors in check by lengthening the G1 phase of the cell cycle

**DOI:** 10.1101/2021.03.03.433829

**Authors:** Emma Coakley-Youngs, Susan Shorter, Medhavi Ranatunga, Simon Richardson, Giulia Getti, Marc Fivaz

## Abstract

CHD8 (Chromodomain Helicase DNA Binding Protein 8) is a chromatin remodeler that preferentially regulates expression of genes implicated in early development of the cerebral cortex. *De novo* mutations (DNMs) in CHD8 are strongly associated with a specific subtype of autism characterized by enlarged foreheads and distinct cranial features. The vast majority of these DNMs are heterozygous loss-of-function mutations with high penetrance for autism. How *CHD8* haploinsufficiency alters the normal developmental trajectory of the human cortex is poorly understood and debated. Previous studies in the mammalian developing cortex have shown progressive lengthening of the G1 phase of the cell cycle as neural stem cells transition from proliferative to neurogenic divisions. G1 length has been proposed to operate as a molecular clock that controls timing of this crucial developmental switch. To determine the influence of CHD8 on cell cycle timing, we disrupted one allele of *CHD8* in human embryonic stem cells (hESCs), differentiated these cells into neural precursor cells (NPCs), and imaged cell cycle progression of individual *CHD8*^*+/−*^ NPCs — in parallel with their isogenic *CHD8*^+/+^ counterparts — during several rounds of cell division. We found a specific and marked decrease in G1 duration in *CHD8*^*+/−*^ NPCs, resulting in an overall shortening of the cell cycle. Consistent with faster progression of *CHD8*^*+/−*^ NPCs through G1 and the G1/S checkpoint, we observed increased expression of E cyclins and elevated phosphorylation of Erk in these mutant cells — two central signalling pathways involved in S phase entry. Together, our findings show dysregulated proliferation of NPCs in a human stem cell model of *CHD8* haploinsufficiency and predict enlargement of the neural progenitor pool in *CHD8*^*+/−*^ developing brains, a phenotype that may explain macrocephaly in individuals with *CHD8* DNMs. Furthermore, our work provides further evidence for a link between autism and cancer and identifies MAPK signaling as a potential therapeutic target for the treatment of this autism subtype.

## Introduction

Recent exome sequencing studies from family trios have led to the identification of numerous spontaneous mutations associated with autism spectrum disorder (ASD) (Barnard et al., 2015; Bernier et al., 2014; Helsmoortel et al., 2014; O’Roak et al., 2012; Weiss et al., 2016; An et al., 2020). Among these, *de novo* mutations (DNMs) in the ATP-dependent chromatin remodeler *CHD8* (Chromodomain Helicase DNA Binding Protein 8) have attracted considerable attention. CHD8 is one of the most frequently mutated gene in ASD — 20 different DNMs have so far been identified, most of which are predicted loss-of-function (LoF) mutations on a single gene copy with high penetrance for ASD (An et al., 2020; Barnard et al., 2015). *CHD8* DNMs give rise to a distinct ASD subtype characterized by enlarged head circumference, dysmorphic facial features and gastrointestinal problems. Notably, macrocephaly is observed in 15% of autistic individuals (Fidler et al., 2000; Fombonne et al., 1999), but the exact nature of this association and the mechanisms underlying macrocephaly in ASD (including the CHD8 ASD subtype), are subject to debate (Durak et al., 2016; Fidler et al., 2000; Gompers et al., 2017).

Chromodomain Helicase DNA-binding proteins belong to one of four subfamilies of chromatin remodelers (Hota and Bruneau, 2016) that use energy from ATP to slide along DNA — a process involved in nucleosome repositioning (Lai and Pugh, 2017). CHD8 expression in the brain peaks during mid-gestational life and is particularly elevated in cortical progenitors and post-mitotic neocortical layers (Bernier et al., 2014), pointing to early corticogenesis as a likely target of *CHD8* DNMs. Accordingly, this chromatin remodeler preferentially influences the expression of genes implicated in neocortical development (Gompers et al., 2017; Suetterlin et al., 2018; Wang et al., 2017), including other ASD risk factors (Cotney et al., 2015; Sugathan et al., 2014). RNA-seq studies in various models of *CHD8* haploinsufficiency identified several gene modules regulated by CHD8: RNA processing, cell cycle regulation, neuronal development/differentiation and synapse function (Gompers et al., 2017; Jung et al., 2018; Katayama et al., 2016; Platt et al., 2017; Suetterlin et al., 2018; Sugathan et al., 2014; Wang et al., 2017), but a clear picture of the role of this chromatin remodeler in brain development is yet to emerge.

While macrocephaly is observed in several independent mouse models of *CHD8* haploinsufficiency (Gompers et al., 2017; Platt et al., 2017; Suetterlin et al., 2018), surprisingly little is known about the function of this gene in cortical assembly. In an early study, silencing of *CHD8* in mouse embryos by RNA interference resulted in reduced proliferation of NPCs and accelerated neuronal differentiation (Durak et al., 2016), an effect attributed to decreased transcription of genes regulating the cell cycle and Wnt signaling. This finding is in line with other reports in non-neuronal tissues describing a stimulatory effect of CHD8 on cell proliferation (Menon et al., 2010; Subtil-Rodríguez et al., 2014) via E2F-dependent transcriptional activation of S phase genes (Subtil-Rodríguez et al., 2014; Rodríguez-Paredes et al., 2009). In marked contrast, germline haploinsufficiency of *CHD8* in mice is associated with a ~ 20% global increase in proliferation of cortical progenitors, and a specific increase in the number of Pax6^+^ radial glial cells located in the subventricular zone of the developing cortex (Gompers et al., 2017). This view is aligned with another recent report describing an inhibitory function of *kismet,* the fly analog of *CHD8*, in intestinal stem cell proliferation (Gervais et al., 2019), and an increasing number of studies implicating *CHD8* as a tumor suppressor (Sawada et al., 2013). While context-dependent functions of CHD8 could account for opposing outcomes in different species and cell types (Rodríguez-Paredes et al., 2009; Wang et al., 2017; Durak et al., 2016), the cause of these conflicting reports in mouse developing cortex is unclear, and is likely to be multifactorial (gene dosage, timing of *CHD8* knockdown by RNAi, cell-autonomous knockdown vs. systemic mono-allelic disruption of *CHD8).* In addition, these two studies are mostly based on pulse-chase incorporation of a nucleotide analog, an approach that can only provide indirect, population-wide information about cell cycling parameters (Nowakowski et al., 1989).

To measure the direct influence of CHD8 on the cell cycle of human neural progenitors, we generaged a stem cell model of *CHD8* haploinsufficiency by CRISPR-Cas9 gene editing and monitored cell cycle progression of individual NPCs with the Fluorescent Ubiquitination-based Cell Cycle Indicator FUCCI (Sakaue-Sawano et al., 2008) by time-lapse microscopy. Using an image analysis pipeline that reconstructs individual cell lineages during several rounds of cell division and measures duration of G1 and S/G2/M during each cycle, we show that heterozygous LoF of *CHD8* results in a marked shortening of G1. Further, we demonstrate that this curtailed G1 phase results from impaired repression of several pathways that drive entry into S phase. Our findings provide strong evidence for an inhibitory function of CHD8 in NPC proliferation and suggest a mechanism for macrocephaly in individuals with *CHD8* DNMs based on abnormal expansion of the progenitor pool in the developing neocortex.

## Material and Methods

### DNA constructs, antibodies and other reagents

The pSpCas9(BB)-2A-Puro (PX459) V2.0 (#62988), pBOB-EF1-FastFUCCI-Puro (#86849), psPax2 (gag-pol) (#12260) and pMD2.G (vsvg) (#12259) DNA plasmids were from addgene. The Pax6 rabbit pAb (Poly19013) was from BioLegend. The OCT4 mouse mAb (3A2A20) was from StemCellTech.The rabbit Phospho-p44/42 MAPK (Erk1/2) (Thr202/Tyr204) mAb (4370) was from CST. All secondary Abs were from ThermoFisher: goat anti-rabbit IgG Alexa FLuor 555 (A27039) and goat anti-mouse IgG Alexa Fluor 488 green (A-11001).

### Maintenance and differentiation of hESCs

The hESCs line H9 (WA09) was from WiCell. We obtained authorization from WiCell (MTA) and from the UK Stem Cell Bank to import and use this line in our laboratory. All stem cell culture reagents were from STEMCELL technologies. We maintained and differentiated hESCs according to STEMCELL protocols.

#### Maintenance

hESCs were grown in mTeSr™ medium on matrigel-coated in (Geltrex A1413201, Thermofisher) tissue culture treated 6 well plates (Corning). Medium was changed daily. hESCs were passaged every 5-7 days using the ReLeSR™ dissociation reagent. Care was taken to passage cells as colonies before they reached confluency. For nucleofection and differentiation, hESCs were dissociated to the single-cell level using the GCDR dissociation reagent and grown in the appropriate medium containing 10 μM Y-27632 (ROCK inhibitor) for the first 24 hrs after dissociation. hESCs were cryopreserved in mFreSR™ and thawed in mTeSr™ containing 10 μM Y-27632 for 12-24hrs until colonies reached a sufficient size.

#### Differentiation

hESCs were dissociated to the single-cell level and grown in STEMdiff™ neural induction media containing dual SMAD inhibitors, in the presence of Y-27632 (10μM) for the first 24 hrs. Medium was changed daily. Cells were passaged a first time after ~ 7 days using accutase, grown for another ~ 7 days in STEMdiff™ (+ dual SMADi) neural induction media and passaged a second time with accutase into STEMdiff™ neural progenitor medium (containing supplements A and B) for final differentiation into NPCs. NPCs were used for experiments after at least one passage in STEMdiff™ neural induction medium. At this point, > 90% NPCs stained positive for Pax6 and negative for Oct4. For all experiments comparing the behavior *CHD8*^*+/+*^ and *CHD8*^*+/−*^ NPCs, cells from these two groups were differentiated in parallel and examined after the exact same number of passages.

### CRISPR-Cas9 gene editing in hESCs

#### sgRNA design, cloning and transfection

20-mer sgRNA were designed using the CRISPR design tool [http://crispr.mit.edu/] against exons 3 and 4 of *CHD8* — the first two conserved exons of the two *CHD8* variants (RefSeq accession numbers NM_001170629.2 and NM_020920.4). We selected two sgRNAs (one against each exon) with the highest score. We then used a plasmid-based procedure for scarless cloning of double-stranded oligonucleotides encoding the sgRNA and its complementary sequence into a cassette containing Cas9 and the sgRNA’s scaffold (Ran et al., 2013). Forward and reverse oligos for each gRNA contained overhangs for directional cloning into the pSpCas9(BB)-2A-Puro plasmid using the BBS1 restriction site. These oligos contained an additional G-C pair upstream of the 20nt gRNA sequence to enhance transcription by the vector’s U6 promotor (Ran et al., 2013). Forward and reverse oligos were phosphorylated with T4 polynuclease kinase, annealed to form dsDNA, and the DNA duplex was cloned into pSpCas9(BB)-2A-Puro by restriction digest with BBS1 and ligation using T4 ligase. Ligation products were then transformed into Sbtl3 competent bacteria. Proper insertion of the sgRNA sequences was confirmed by Sanger sequencing. gRNA-Cas9 constructs were then separately introduced in hESCs using the Amaxa 4D-nucleofector (Lonza) and the nucleofection program/kit designed for the H9 hESC line. For this, hESCs were dissociated at the single cell level and deposited in a 16-well Nucleocuvette strip in presence of the ROCK inhibitor Y27632 (200,000 cells per well) and 2 μg of ex3-gRNA-Cas9 or ex4-gRNA-Cas9 or control plasmids containing the Cas9-expressing vector without sgRNA or a GFP-expressing plasmid to evaluate transfection efficiency. We typically obtained 60-70% transfection efficiency using this protocol. Cells were immediately transferred to a 24-well plate after electroporation and grown for another 24hr in presence of Y27632.

#### Surveyor assay

Indel frequency was assessed using the surveyor assay (Integrated DNA Technologies, IDT) according to the manufacturer’s instructions. ~ 500bp genomic DNA fragments encompassing the predicted DSB in both exon 3 and 4 were amplified from nucleofected hESCs using primers described in Table 1. These genomic amplicons were then denatured, re-annealed and digested with the mismatch-specific surveyor endonuclease. The digestion patterns we obtained from hESCs transfected with ex3-gRNA-Cas9 and ex4-gRNA-Cas9 constructs are consistent with the presence of indels at the site predicted by the sgRNA sequence (Figure S1a in Supplemental Data). These digested PCR products were not observed in hESCs transfected with a GFP plasmid (Figure S1a).

**Table 1.**
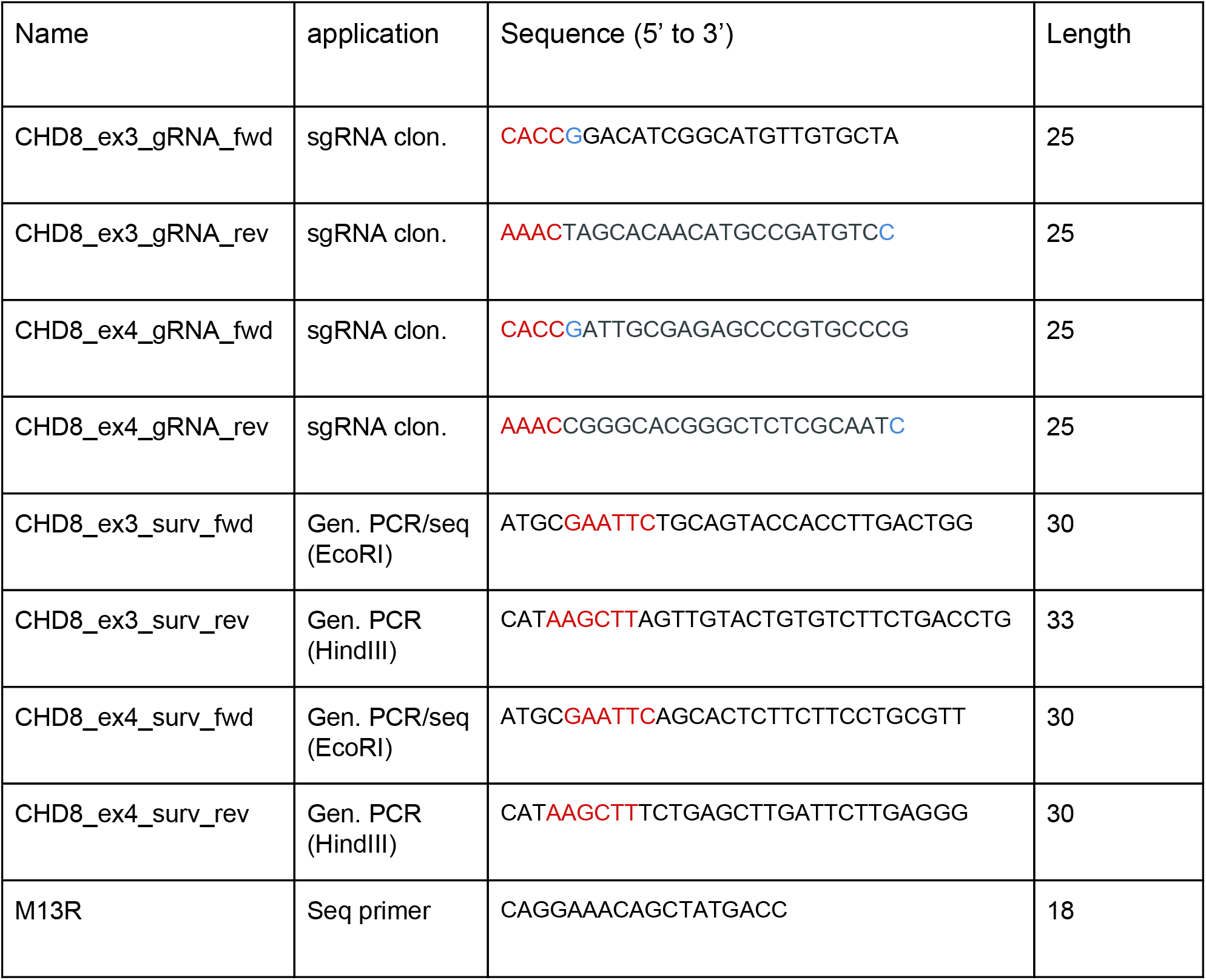
DNA oligos for sgRNA cloning, genomic PCR and single-allele sequencing.

#### Isolation of single CRISPR clones

After confirmation of successful editing by the surveyor assay, we isolated individual clones of hESCs by limited dilution in matrigel-coated 10cm dishes (56.7 cm^2^). Individual colonies were manually picked under the microscope and seeded in a 96 well plate for expansion and cryopreservation. For each sgRNA, about 50 clones were successfully expanded and screened for the presence of indels by DNA sequencing of a genomic amplicon. We used the same primers for genomic PCR and sequencing of a ~500 bp region containing the predicted edited site in exon 3 and 4 (Table 1).

#### TIDE analysis and single-allele sequencing

Genomic amplicons originating from individual hESC clones were further analyzed by the TIDE software (Brinkman et al., 2014) for deconvolution of mixed sequences and prediction of indel type and frequency (www.tide.nki.nl). For single allele sequencing, genomic amplicions (amplified with primers containing the HINDIII and ECOR1 sites, Table 1) were cloned by restriction digest into the pUC19 plasmid and sequenced using the M13R oligo (Table 1). More than 30 transformants were sequenced to obtain a reasonable estimate of mutation frequency.

### TaqMan RT-qPCR

Total RNA was isolated from hESCs or NPCs grown in a 6-well plate (9.5 cm^2^) using the RNeasy minikit (Qiagen). A DNase 1 digestion step was performed during RNA extraction to degrade remaining genomic DNA. 2μg total RNA was reverse-transcribed into cDNA using the high capacity cDNA reverse transcription kit (Thermofisher, 4368813). RT-qPCR was performed with Taqman assay chemistry in a 96 well format using a QuantStudio1 RT-qPCR machine (Applied Biosystems). TaqMan probes (Table 2) were pre-designed by ThermoFisher across exon-exon boundaries. Fold-change in transcript expression was calculated using the 2^−ΔΔCt^ method. ΔCt was calculated by subtracting the Ct value of the control gene POLR2a from that of the target gene (probe). ΔΔCt was calculated by subtracting the ΔCt of the target gene (probe) in CHD8^+/+^ (wildtype) cells from that in CHD8^+/−^ (mutant) cells. Fold-change in gene expression is the average of 3 independent experiments (n = 3). For each independent experiment, Ct was measured in triplicates or quadruplicates.

**Table 2.**
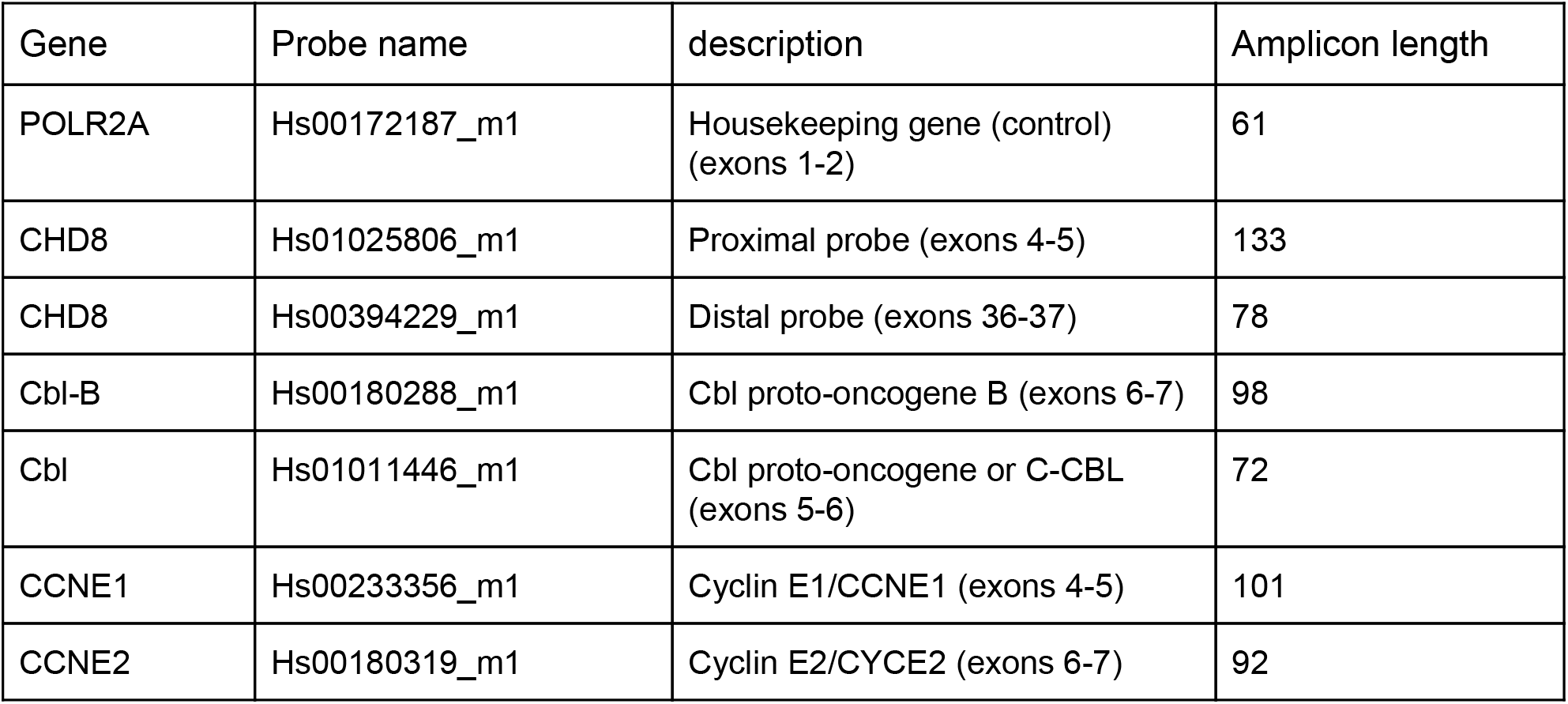
RT-qPCR TaqMan probes.

### Flow cytometry

The distribution of hESCs and NPCs in the different phases of the cell cycle was measured by DNA content analysis using flow cytometry. hESCs or NPCs were cultured in matrigel-coated 6-well plates (9.5 cm^2^) until they reached ~70% confluency, dissociated into a single-cell suspension, fixed with EtOH and stained with propidium Iodide (ab139418, abcam) according to the manufacturer’s instructions. *CHD8*^*+/+*^ and *CHD8*^*+/−*^ cells were then analyzed in parallel on a BD Accuri C6 flow cytometer. Cell debris and doublets were gated out using forward and side scatter density plots (FSC-A vs SSC-A) and forward scatter height versus forward scatter density area plots (FSC-A vs FSC-H) respectively. PI fluorescence was detected using the PE-A channel. PI histograms (cell count vs PI intensity) were then generated and gated to isolate G0-G1, S and G2-M phases (Figure 2c). The % of cells in each of these three groups was then computed and averaged from 3 independent experiments comparing the two genotypes in hESCs and NPCs.

### Lentiviral production

Low passage Hek293FT cells were grown in 5 × T175 flasks in G1 medium (DMEM supplemented with 10% FBS, 1 mM sodium pyruvate, G418 and PenStrep). When cells reach 70-80% confluency, they were transfected with the following three DNA plasmids: pBOB-EF1-FastFUCCI-Puro, psPax2 (gag-pol) and pMD2.G (vsvg), at a 3:2:1 ratio using Lipofectamine 3000. 6 hours post transfection cells were switched to Ultraculture medium (Lonza) and grown for another 24-48hrs. The supernatant containing the viral particles was collected, cleared by low-speed centrifugation (1000rpm for 5 min) and filtered on a 0.45 μm filter-flask. The filtered supernatant (~ 120 ml) was then placed at the bottom of two conical centrifuge tubes, overlaid with 20 ml of 20% sucrose in PBS and spun at 20’000g for 2 hrs at 4°C. The viral pellet was then resuspended in 1.5 ml ice-cold PBS from each centrifuge tube. Lentiviral particles were cleared by a final spin at 1000g for 10 minutes (4°C). The supernatant was alicoted (5 μL) and immediately frozen at −80°C.

### Live-cell confocal imaging and analysis of cell cycle timing

Live-cell imaging was carried out in a temperature (37°C) and CO_2_-controlled environment using a Micro Confocal high-content imaging system (Molecular Devices, USA) or laser-scanning confocal LSM 880 with airy scan (Zeiss, Germany).

#### Analysis of cell cycle timing

To extract cell cycle timing information from FUCCI time series, we developed a MATLAB-based interactive point-and-click segmentation algorithm. A binary image stack (time series) is first generated for each channel (green: mAG-Geminin_1-110_, red: mKO2-Cdt1_30-120_) using an adaptive thresholding approach. Color-specific binary stacks are then combined into a single stack which is used for click-based segmentation. Point-and-click is performed by the user on a merged RGB image of the green and red channels containing an interactive cursor to facilitate identification of cells. A cell and its progeny are then identified by a mouse click iteratively through the time series using the “bwselect” function in matlab. The sequence of mouse clicks defines cell identity throughout the stacks allowing to build cell lineages. The mean green and red fluorescence intensity in each segmented/tracked nucleus is then recorded and plotted as a function of time. Cell cycle phase duration (G1 and S/G2/M) is then computed based on the green and red intensity profiles for each cell in a lineage. The matlab script required to run this analysis is available in Supplemental Material.

### Immunofluorescence

Cells were seeded on matrigel-coated glass coverslips, fixed with 4% paraformaldehyde, 4% sucrose for 20 minutes, permeabilized with 0.5% Triton-X-100 and blocked with 10% HSA (H0146) and 1% BSA (15561020, Thermofisher) in 1XPBS. Cells were labelled with primary antibody 1:200 in 1% BSA, then secondaries 1:1000 in 1% BSA in the dark. Cells were mounted with Mowiol (Sigma). Imaging was performed with an LSM 880 confocal microscope, (Zeiss, Germany).

### Statistical analysis

Boxplots and swarmplots were generated with Python (Jupyter notebook) using the seaborn data visualization library. The box shows the quartiles (25% and 75%) of the dataset. The whiskers show the rest of the distribution contained within 1.5x the IQR (interquartile range). The median of the distribution is indicated by a horizontal bar. We used the Python SciPy library for statistical analysis. The nonparametric Mann-Withney U test was employed to compare cell cycle timing data and pErk expression in *CHD8*^+/+^ and *CHD8*^+/−^ cells. P values were corrected for multiple hypothesis testing using the post-hoc Bonferroni method.

## Results

### Mono-allelic disruption of *CHD8* in hESCs by CRISPR-Cas9 gene editing

The human CHD8 gene encodes two transcripts with a different 5’ exonic structure (Figure 1a). We designed two small guide RNAs (sgRNAs) against the first two conserved exons (Figure 1a), cloned them into a gRNA-Cas9 expressing DNA plasmid and nucleofected these gRNA-Cas9 constructs into hESCs. After confirming successful editing of the *CHD8* locus using the surveyor assay (Figure S1a in Supplemental Material), we isolated individual clones by limited dilution (about 50 clones for each sgRNA) and sequenced a ~500bp genomic amplicon containing the predicted edited sites. We obtained 4 to 6 edited clones per sgRNA indicating an editing frequency of ~10%. To identify clones with a putative LoF indel, we analyzed DNA sequences displaying a clear breakpoint (Figure 1b) with the TIDE (Tracking of Indels by Decomposition) software (Brinkman et al., 2014). TIDE identified several clones with a predicted mono-allelic indel (not shown). Based on TIDE’s analysis, we selected clone E11 for further studies. For E11, TIDE predicted a heterozygous 1bp insertion in exon 4 at the predicted double strand break (Figure S1b in Supplemental Data). We confirmed TIDE’s results by single-allele sequencing of E11 (Figure 1c). Out of the 34 alleles we sequenced, 15 were wt and 19 contained the 1bp insertion (Figure 1d), consistent with a heterozygous mutation. The frameshift induced by this 1bp insertion resulted in the introduction of a stop codon 8bp downstream of the mutation (Figure 1e). Next, we quantified *CHD8* transcripts levels in wt and E11 hESCs using RT-qPCR. Two sets of TaqMan probes were used (Figure 1a) annealing across exon-exon boundaries shortly after the DSB (exon 4-5, proximal probe) or towards the end of the transcript (exon 36-37, distal probe). We measured a decrease in the *CHD8* message of 49 ± 0.06% (std) and 34 ± 0.15% (std) in E11 with the proximal and distal primer sets respectively (Figure 1F). Collectively, these results show successful disruption of one copy of the *CHD8* gene in the E11 hESC line, with no apparent compensation from the remaining wt allele. Finally, the E11 clone expressed the pluripotent stem cell marker Oct4 in the vast majority of cells, similar to wt hESCs (Figure 2a,b) indicating that mono-allelic disruption of *CHD8* has not compromised the pluripotent state of these cells. We therefore used the E11 line as a stem cell model of *CHD8* haploinsufficiency in subsequent experiments.

**Figure 1.**
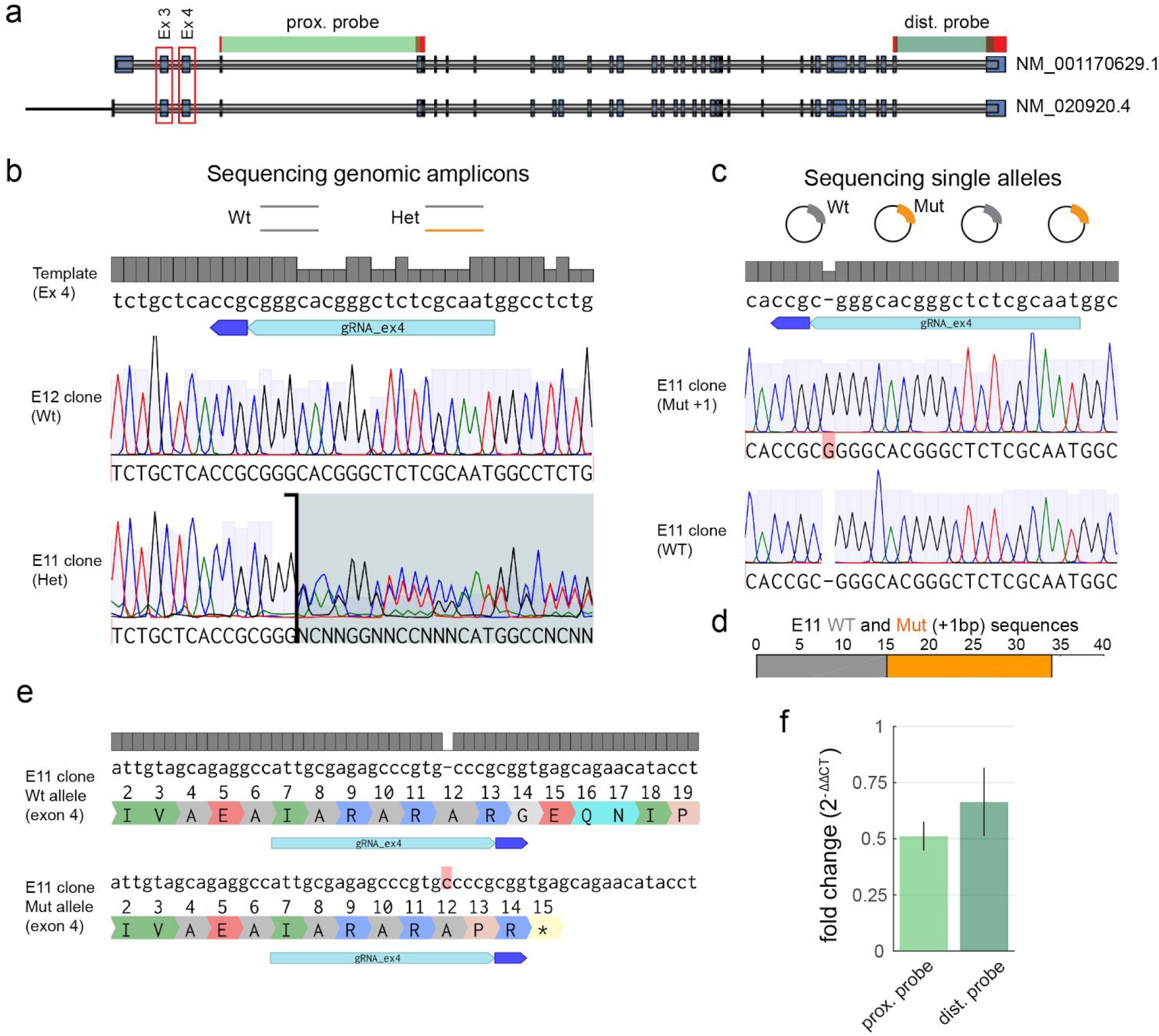
Disruption of a single allele of *CHD8* in hESCs by CRISPR-Cas9 gene editing. (a) Genomic organization of the *CHD8* locus. Boxed in red are the first two conserved exons of the two *CHD8* variants against which sgRNAs were designed. The regions across exon-exon boundaries amplified by proximal and distal TaqMan primers are indicated in green. (b) Genomic sequencing of two individual clones — E12 and E11 — isolated from H9 cells transfected with a gRNA-Cas9 construct targeting exon 4. E11 shows a clear breakpoint at the Cas9 predicted cleavage site resulting from mixed allelic DNA sequences. (c) Single-allele sequencing reveals the insertion of a single nucleotide C (G in the opposite strand sequenced here) at a frequency consistent with a heterozygous mutation (d). (e) DNA and protein sequence alignments of the wt (top) and mutated (bottom) allele of the E11 clone. The one base pair insertion introduces a stop codon shortly after the edited site. (f) TaqMan RT-qPCR showing fold-change in the *CHD8* mRNA transcript in E11 relative to wt hESCs (n = 3). Error bars indicate std.

**Figure 2.**
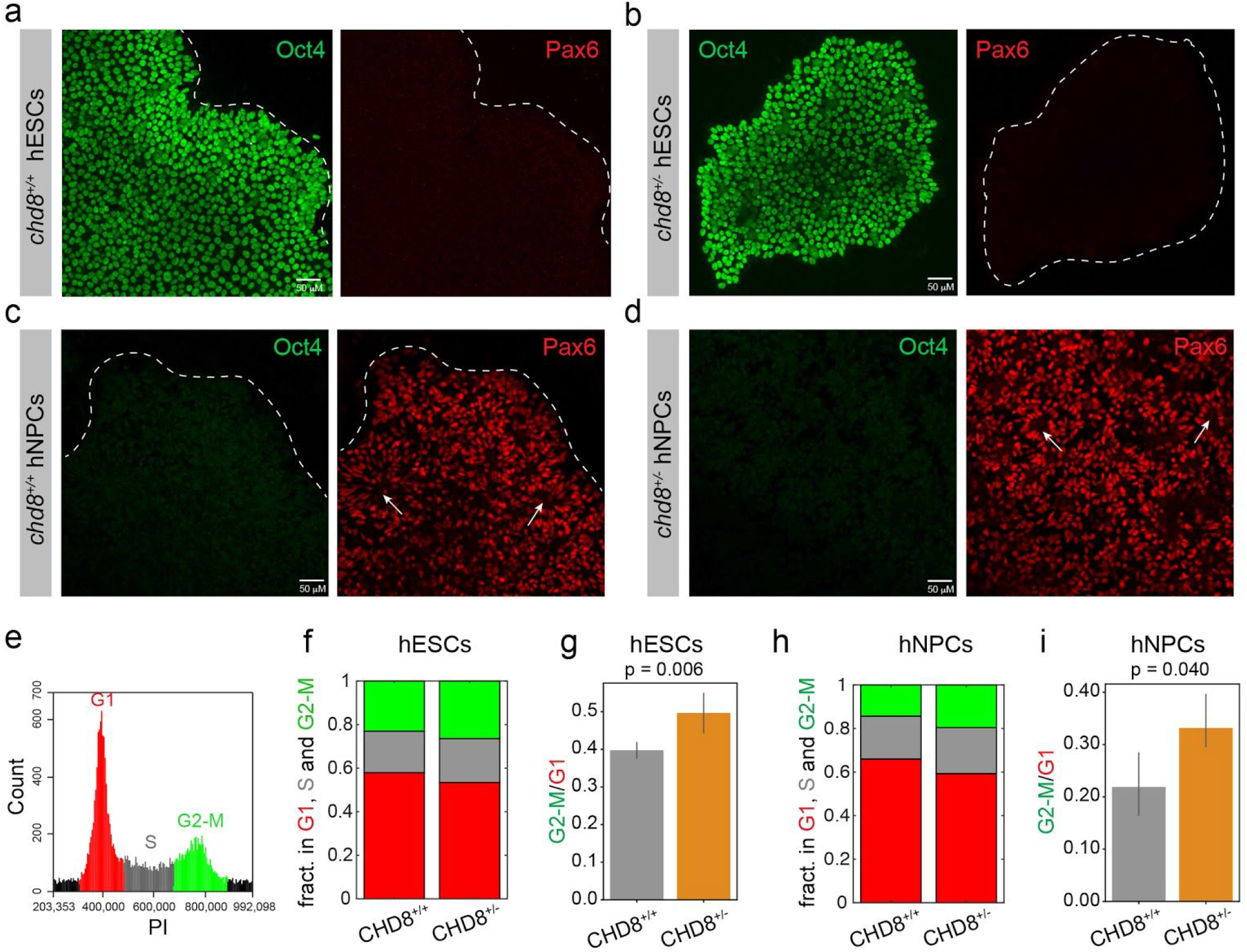
Flow cytometry reveals subtle differences in cell cycle profile of *CHD8*^+/−^ hESCs and hNPCs. (a-d) Immunostaining of Oct4 (green) and Pax6 (red) in *CHD8*^+/+^ (a,c) and *CHD8*^+/−^ (b,d) hESCs (a,b) and hNPCs (b,d). Arrows point to neural rosettes. (e-i) Cell cycle profile by DNA content analysis. (e) PI Histogram (cell number vs PI intensity) showing the distribution of cells in G0-G1, S and G2-M. (f,h) fraction of wt and mutant cells in G0-G1, S and G2/M for hESCs (f) and hNPCs (h). (g,i) Mean G2-M/G1-G0 ratio for hESCs (g) and hNPCs (i). n=3, error bars indicate std. p values were obtained by a Mann-Withney U test.

### Heterozygous disruption of *CHD8* increases the proportion of cells in S/G2/M at the expense of G0-G1

hESCs were next differentiated into NPCs using a neural induction procedure based on dual-SMAD inhibition (Chambers et al., 2009; Gerrard et al., 2005; Muratore et al., 2014).This protocol efficiently converts hESCs into NPCs of the dorsal telencephalon (forebrain) lineage which progressively acquire the features of radial glial cells — the main source of neural stem cells in the developing cortex. We confirmed efficient differentiation of hESCs into NPCs by the concomitant loss of the pluripotent stem cell marker Oct4 and acquisition of the cortical progenitor marker Pax6 (Figure 2a-d). Neural induction was quasi-complete for both *CHD8*^*+/+*^ and *CHD8*^*+/−*^ (E11) lines and led to the spontaneous assembly of neural rosettes (Figures 2c,d) — a defining behaviour of cortical progenitor cells (Ziv et al., 2015). To determine whether CHD8 influences the cell cycle, we first measured the distribution of cells in G0-G1, S and G2-M by flow cytometry (Figure 2e-i). In hESCs, heterozygous disruption of *CHD8* led to a modest decrease in the fraction of cells in G0-G1 and a corresponding increase in the number of cells in S and G2-M (Figure 2f,g). A similar but more pronounced effect was observed in NPCs (Figure 2h,i). As expected, a higher fraction of NPCs were in G1 compared to hESCs (Figure 2f,h) consistent with a longer G1 phase in neural progenitors (Calegari et al., 2005; Lange et al., 2009). These data show that that a higher proportion of *CHD8*^*+/−*^ NPCs and, to a lesser extent, *CHD8*^*+/−*^ hESCs, are in S and G2-M, suggesting that these mutant cells proliferate faster.

### Imaging cell cycle progression reveals marked shortening of G1 phase in *CHD8*^*+/−*^ NPCs

To directly measure duration of cell cycle phases, we imaged cell cycle progression using FUCCI (Sakaue-Sawano et al., 2008) by dual-color live-cell confocal microscopy. This probe labels G1 phase nuclei in red and those in S/G2/M phases in green (Figure 3a-c) without interfering with cell cycle dynamics (Pauklin and Vallier, 2013). To faithfully compare cell cycle progression in *CHD8* wt and mutant cells, both groups were simultaneously imaged in multiple wells of a 96 well plate for 20 to 70 hrs, by high-content microscopy. FUCCI was introduced in these cells by lentiviral delivery using a viral titer that led to 10-20% of FUCCI-expressing cells to allow tracking of individual cell lineages (Figure S2a in Supplemental Data). Cells remained healthy and proliferative throughout the imaging session, as judged by a clear increase in colony size with time (Figure S2b,c in Supplemental Data). The size of these data sets precluded manual analysis of cell cycle timing. Full automatization of cell tracking is difficult, however, because during conversion from green to red (M-G1), FUCCI emits no or little light for several successive frames (Figure 3c-e. We thus wrote an interactive image processing algorithm that segments a cell and its progeny based on mouse clicks and plots dual-color fluorescent traces for mother and daughter cells (see methods). G1 and S/G2/M lengths were then computed based on these time series (Figure 3c). We estimate that this computer-assisted approach speeds up the analysis by a factor ~ 10.

**Figure 3.**
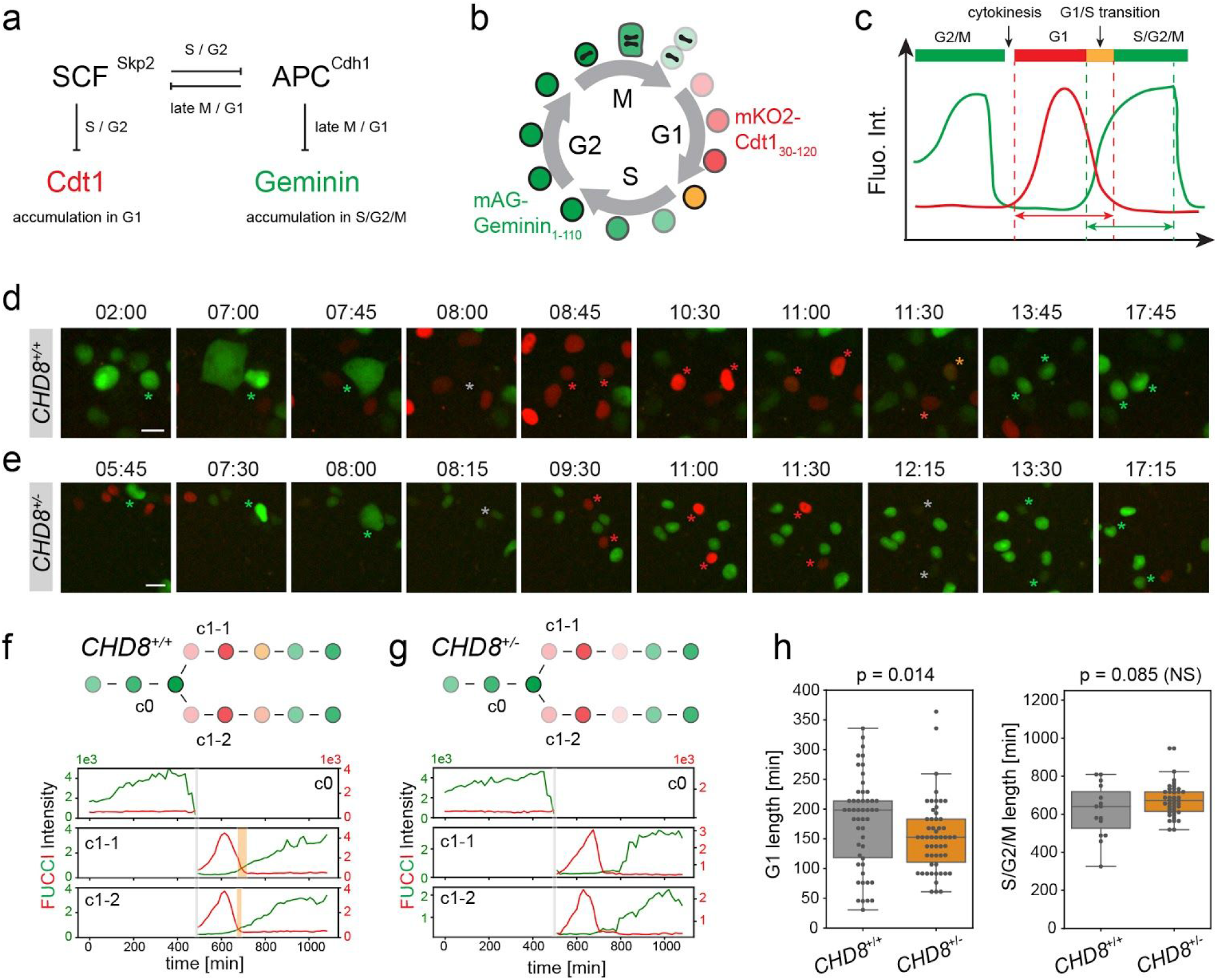
Cell cycle timing in hESCs. (a,b) Color-based detection of G1 and S/G2/M by the FUCCI fluorescent reporter. The G1 and S/G2/M probes are expressed as a single open reading frame using the T2A self-cleaving peptide sequence ensuring stochiometric expression of both probes. (c) Measurement of G1 and S/G2/M duration based on FUCCI intensity traces (see methods). (d-h) High-content FUCCI imaging in wt (d,f,h) and mutant (e,g,h) hESCs. (d,e) Snapshots of *CHD8*^+/+^ (d) and *CHD8*^+/−^ (e) hESCs during cell cycle progression (~ 17 hrs). Asterisks show a cell initially in S/G2 undergoing mitosis and cytokinesis and its two daughter cells transitioning from G1 to S/G2. Time is in hours and mins. Scale is 20 μm. (f,g) FUCCI intensity traces of mother and daughter cells marked by an asterisk in (d,e). The green y axis (left) indicates mAG-Geminin_1-110_ intensity (S/G2/M), the red y axis (right) indicates mKO2-Cdt1_30-120_ intensity (G1). Grey bars indicate cytokinesis (M/G2). Orange bars indicate G1/S transitions. (h) Box plots showing duration of G1 and S/G2/M in *CHD8*^+/+^ (n = 48) and *CHD8*^+/−^ (n = 49) cells. p values were measured using a Mann-Withney U test.

Using this image analysis pipeline, we first examined cell cycle timing in hESCs. *CHD8*^*+/+*^ and *CHD8*^*+/−*^ hESCs went through G1 in less than 3 hrs while they spent at least 10 hrs in S/G2/M (Figure 3d-h). A short G1 phase with no apparent regulation of the G1 checkpoint is a hallmark of embryonic pluripotent stem cells (Neganova et al., 2009; Pauklin and Vallier, 2013; Neganova et al., 2009, 2011; Pauklin and Vallier, 2013). Accordingly, we rarely captured a clear G1/S transition in these cells (regardless of genotype), as reflected by little or no overlap between mKO2-Cdt1_30-120_ and mAG-Geminin_1-110_ expression (Figure 3c). Occasionally, transient co-expression of mKO2-Cdt1_30-120_ and mAG-Geminin_1-110_ could be detected in wt but not in mutant cells (Figure 3d-g). Notably, however, G1 length was shorter in *CHD8*^*+/−*^ cells compared to their *CHD8*^*+/+*^ counterparts — median ± std of 153 ± 61 min vs 198 ± 78 min (p = 0.014) — (Figure 3h), indicating that this phase of the cell cycle is further truncated in mutant cells. No significant difference in S/G2/M timing was observed between the two genotypes, although the relatively short duration of this live-cell experiment (20 hrs) precluded a precise comparison of S/G2/M lengths.

Next, we imaged cell cycle progression in stem-cell derived neural progenitors for up to three days (70 hrs). We were able to track individual cell lineages in both wt and mutant hNPCs over several rounds of cell division (Figure 4a-d). NPCs were more motile than their undifferentiated precursors, displayed a characteristic boomerang-shaped nucleus (Figure 4a,b) and, as expected, progressed more slowly through the cell cycle (Figure 4e). G1 in particular was significantly longer in hNPCs than in hESCs — up to 10x in wt cells (compare Figures 4e and 3h). G1/S transitions (overlapping green and red FUCCI signals) were also protracted in hNPCs consistent with the presence of a proper G1/S checkpoint in these cells. Comparative analysis of phase duration across genotypes revealed a marked shortening of G1 in *CHD8*^*+/−*^ hNPCs compared to their wt counterparts — median ± std of 517 ± 410 min vs 1408 ± 982 min (p = 7 × 10^−7^ (Figure 4c-e). In contrast, no overt change in S/G2/M duration was observed in mutant cells (Figure 4c-e) indicating a profound and specific effect of *CHD8* haploinsufficiency on the G1 phase of the cell cycle. Collectively, these imaging experiments show that disruption of one *CHD8* copy is sufficient to severely shorten G1 and facilitate the G1/S transition in hNPCs, an effect that likely boosts proliferation of neural progenitors and delays their differentiation into neurons. The flip side of these results is that CHD8 represses proliferation and self-renewal of hNPCs.

**Figure 4.**
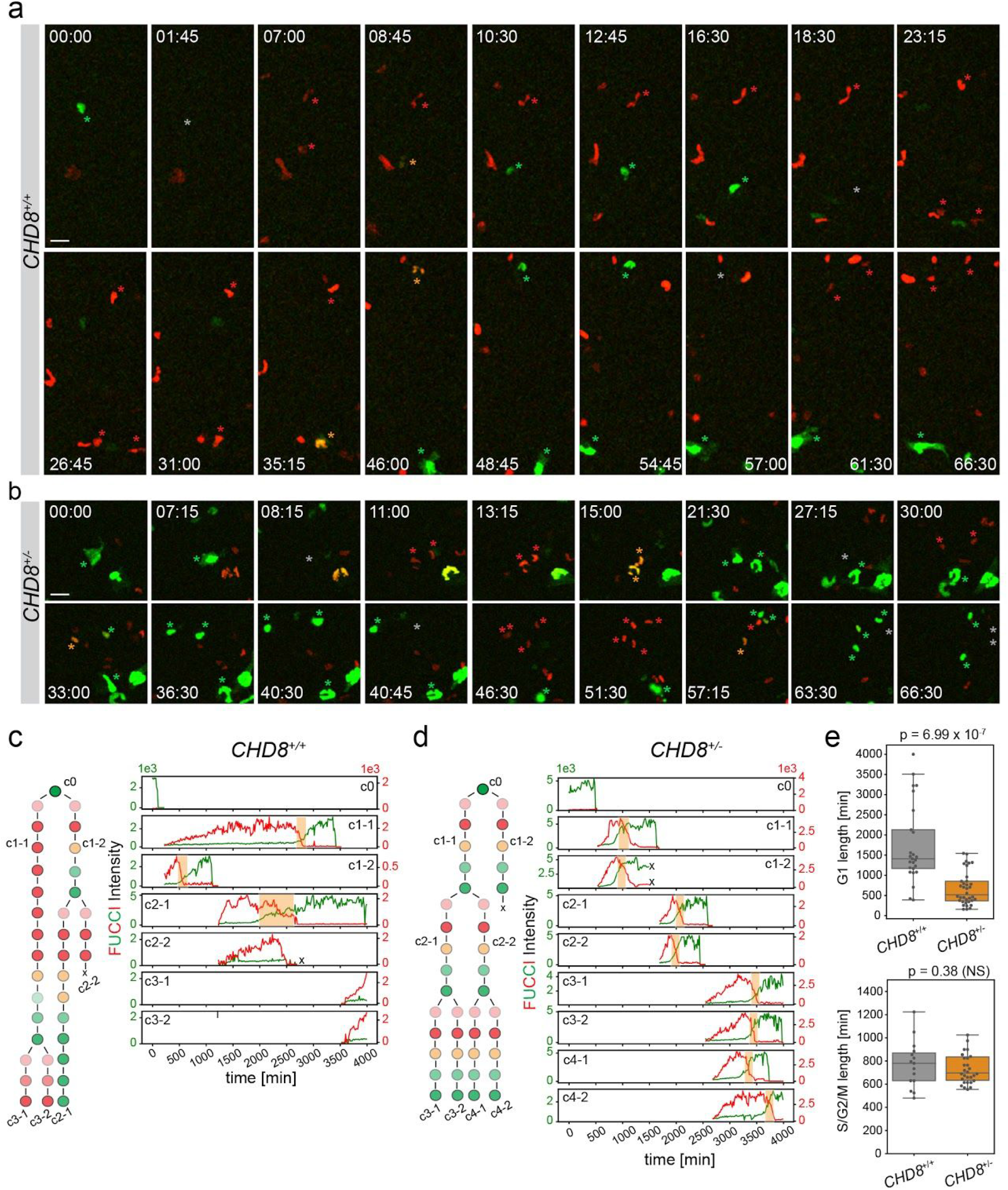
Shortened G1 length in *CHD8*^+/−^ hNPCs. (a,b) Time-lapse imaging of clonal lineages in wt (a) and mutant (b) NPCs going through several rounds of cell division over the course of ~70 hrs. Cells corresponding to a single lineage are marked by asterisks which are color-coded according to cell cycle phase. Time is indicated in hours and mins. Scale bar: 20 μm. (c,d) Lineage reconstruction and FUCCI intensity traces of individual cells marked by an asterisk in (a,b) for wt (c) and mutant (d) NPCs. The green y axis (left) indicates mAG-Geminin_1-110_ intensity (S/G2/M), the red y axis (right) indicates mKO2-Cdt1_30-120_ intensity (G1). Orange bars shown G1/S transitions. (e) Box plots showing duration of G1 and S/G2/M in *CHD8*^+/+^ (n = 23) and *CHD8*^+/−^ (n = 34) cells. p values were measured using a Mann-Withney U test.

### CHD8 represses pathways regulating S phase entry

To explore the mechanisms by which CHD8 lengthens G1, we searched for CHD8 target genes that regulate G1 and the G1/S checkpoint. Cyclins E1 and E2 are expressed during the G1/S transition and have been proposed to drive S phase entry. The expression of both E cyclins is modulated by CHD8, but the directionality of this regulation appears to be context-dependent (Rodríguez-Paredes et al., 2009; Shingleton and Hemann, 2015). A comparative RT-qPCR analysis showed that cyclin E1 (CCNE1), and to a lesser extent cyclin E2 (CCNE2), are upregulated in *CHD8*^*+/−*^ hNPCs, while as expected, *CHD8* transcripts levels were halved (Figure 5a). Elevated expression of cyclins E1 and E2 in mutant hNPCs is consistent with faster progression of these cells through the G1/S checkpoint and, thus, shorter G1. EGF-family growth factors and downstream MAPK activation is another key pro-mitotic signalling cascade regulating S phase entry (Kobayashi et al., 1998). Recently, *kismet*, the fly ortholog of mammalian *CHD7* and *CHD8,* was shown to downregulate the EGF receptor (EGFR) by upregulating transcription of the E3 ligase Cbl known to promote degradation of the EGFR (Gervais et al., 2019). To determine whether CHD8 transcriptionally regulates EGF signaling in neural progenitors, we first compared expression of Cbl in *CHD8*^*+/−*^ and *CHD8*^*+/+*^ hNPCs. In mammalian cells, two closely-related Cbl isoforms — Cbl and Cbl-b — have been implicated in EGFR degradation (Mohapatra et al., 2013). While Cbl expression is not influenced by CHD8, Cbl-b transcripts levels were 20% down in *CHD8*^*+/−*^ hNPCs (Figure 5a) indicating a mild repressive activity of CHD8. We next examined the impact of CHD8 on MAPK signalling. Immunostaining against a phosphorylated form of ERK1/2 (Thr202/Tyr204) revealed a marked increase in the activity of this MAPK in *CHD8*^*+/−*^ hNPCs, particularly in the nucleus (Figure 5b-f). Together, these results show that CHD8 represses two central signaling pathways that regulate progression through G1. Disruption of one *CHD8* allele is sufficient to relieve this transcriptional inhibitory circuit and accelerate S phase entry in neural progenitors (Figure 6).

**Figure 5.**
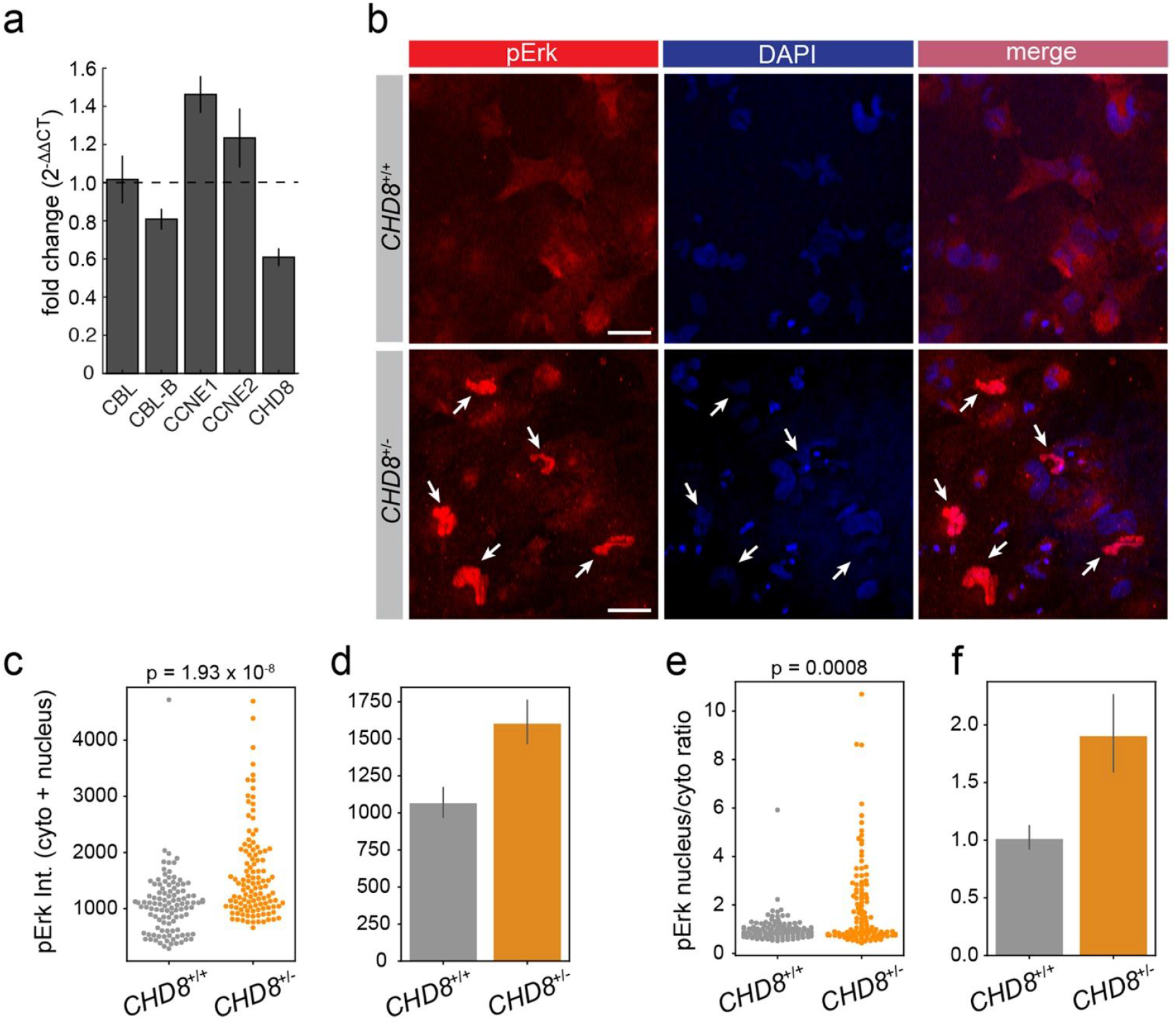
CHD8 represses cyclins E and MAPK signalling. (a) RT-qPCR analysis of *CHD8*^+/+^ and *CHD8*^+/−^ hNPCs. Fold-change (mut/wt) in mRNA expression is shown for the indicated genes (n=4). (b) pErk immunostaining in *CHD8*^+/+^ and *CHD8*^+/−^ hNPCs. Arrows point to pERK in the nucleus. Scale is 20 μm. (c-f) Quantification of pERK total intensity (c,d) and nucleus to cytoplasm intensity ratio (e,f) in *CHD8* wt (n = 109) and mutant cells (n = 116). (c,e) swarmplots. Bar graphs (d,f) show mean Intensity ± 95% CI (confidence interval). p values were measured using a Mann-Withney U test.

**Figure 6.**
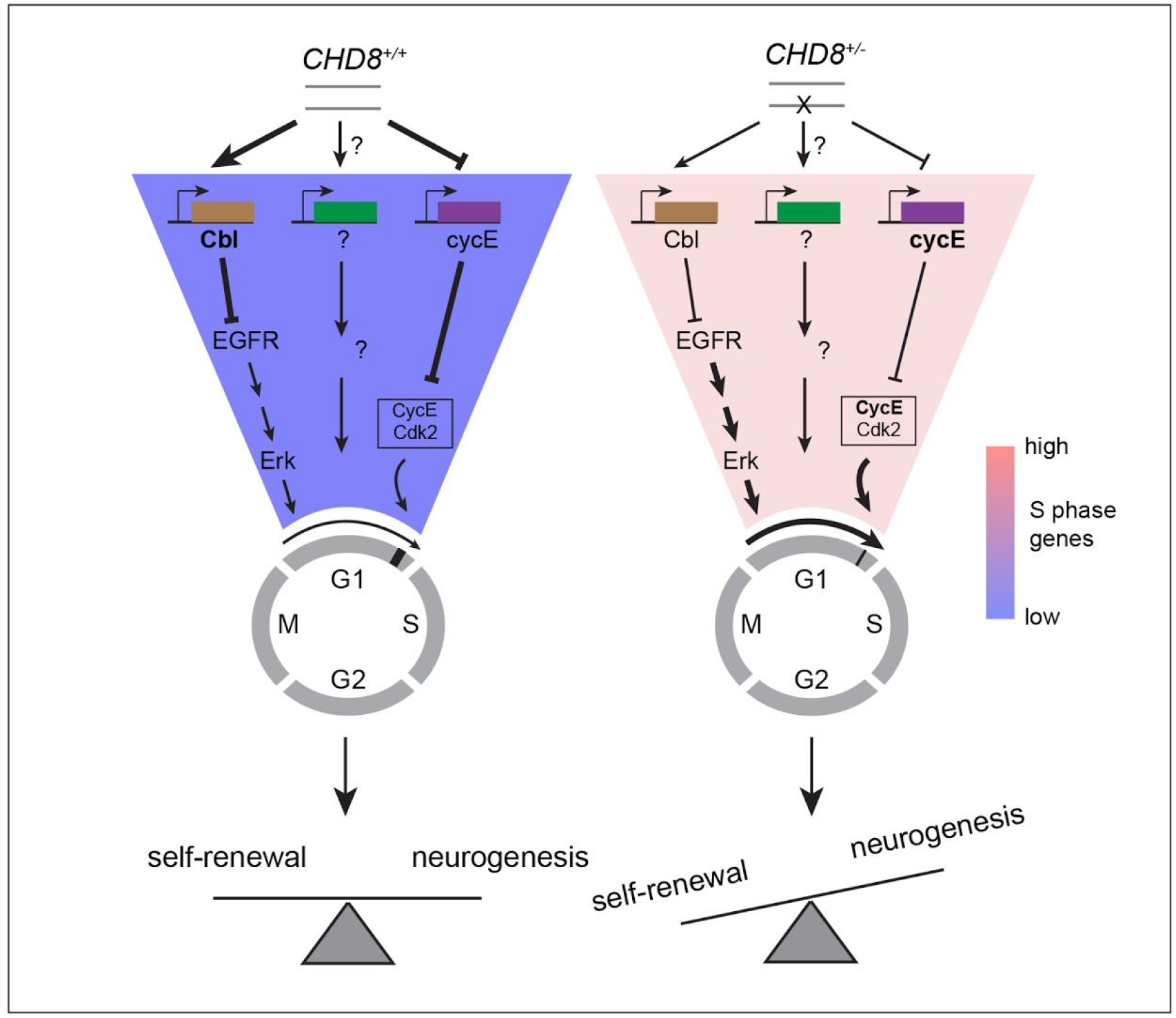
A model for CHD8 regulation of G1 length and S phase entry. Loss-of-function of a single *CHD8* allele shortens the G1 phase of the cell cycle in neural stem cells by relieving transcriptional repression of the MAPK pathway and cyclins E. Truncated G1 causes overproliferation of cortical progenitors by accelerating the cell cycle and by promoting self-renewing divisions at the expense of neurogenic ones.

## Discussion

Cell cycle timing is of central importance for organ growth and maintenance. Changes in duration of cell cycle phases, G1 in particular, have been proposed to control the balance between stem cell proliferation and differentiation and are thought to play instructive roles in cell fate decisions at different stages of development — from the original differentiation of ESCs into the three primary germ layers (Pauklin and Vallier, 2013) to the final assembly of organs including the brain (Dehay and Kennedy, 2007). ESCs and PSCs typically have a short G1 phase which reflects the lack of a proper G1/S checkpoint. Differentiation of stem cells is invariably associated with lengthening of G1, a process thought to broaden the receptive window of these cells to extrinsic cues. Increase in G1 duration is also observed in the mammalian developing cortex, as progenitors transition from proliferative to neurogenic modes of cell division (Dehay and Kennedy, 2007; Lukaszewicz et al., 2002; Calegari et al., 2005; Lange et al., 2009).

In agreement with these findings, we observed a pronounced lengthening of G1, up to 10-fold in wt cells, after differentiation of hESCs into hNPCs, with no overt change in other phases of the cell cycle (Figures 3 and 4). The median G1 duration in wt hNPCs is particularly long (29 hrs) and is characterized by a large variance across individual cells (Figure 4e), possibly reflecting cell cycle exit and re-entry events (Dehay and Kennedy, 2007; Pajalunga et al., 2008). G1 length only increases about 3x in *CHD8*^*+/−*^ NPCs relative to their undifferentiated precursors and is also about 3x shorter than that of their *CHD8*^*+/+*^ counterparts. Given that the median difference in G1 between wt and mutant NPCs is ~ 15hrs (900 min), this truncated G1 phase results in more than a 2-fold shortening of the cell cycle in *CHD8*^*+/−*^ NPCs. Mathematical simulations have shown that halving the length of the cell cycle doubles the rate of neuron production (Caviness et al., 2003), a straight-forward effect that predicts a substantial increase in the number of cortical neurons in the adult brain. A 3-fold shortening of G1 in progenitors is also expected to severely affect the balance between proliferation and differentiation. G1 shortening by overexpression of Cdk4/cyclinD1 promotes expansion of mouse progenitors and delays neurogenesis (Lange et al., 2009). This, together with other reports indicating shortening of G1 in cortical progenitors exposed to mitotic cues and, conversely, increase in G1 length by differentiation factors (Lukaszewicz et al., 2002), suggest that cell cycle timing may influence the switch from self-renewing to neurogenic divisions. Based on these data, it is tempting to speculate that shortening of G1 in CHD8^+/−^ NPCs promotes proliferative divisions at the expense of neurogenic ones thus further contributing to the expansion of the neural progenitor pool. This model also predicts delayed neuronal differentiation and maturation in *CHD8*^*+/−*^ developing cortices, a form of neoteny recently described in both mouse and iPSC models of *CHD8* haploinsufficiency (Gompers et al., 2017; Liu et al., 2019), and other animal models of autism (Chomiak and Hu, 2013). Together, our findings support the notion that macrocephaly in the CHD8 ASD subtype (and in animal models of CHD8 haploinsufficiency) results primarily from an accelerated expansion of the neural progenitor pool.

How does CHD8 prolong the G1 phase of the cell cycle ? We provide evidence that CHD8 represses two key pathways that regulate progression through G1 and S phase entry. Disruption of one copy of *CHD8* results in transcriptional upregulation of E cyclins, cyclin E1 in particular. E cyclins are turned on in late G1 and together with their cognate cyclin-dependent kinase cdk2 regulate G1/S checkpoint transition in ESCs (Neganova et al., 2009, 2011) and cortical progenitors (Delalle et al., 1999; Lukaszewicz et al., 2005; Watanabe et al., 2015). This inhibitory effect of CHD8 on E cyclins expression is in good agreement with a recent RNAseq study in *CHD8*^*+/−*^ iPS-derived cerebral organoids (Wang et al., 2017) but contrasts with an earlier report in HeLa cells showing an opposite effect of CHD8 on cyclin E transcription (Rodríguez-Paredes et al., 2009). These seemingly contradictory results probably reflect context-dependent transcriptional regulation by CHD8 (Rodríguez-Paredes et al., 2009; Wang et al., 2017; Durak et al., 2016), a feature particularly apparent in cancer, where CHD8 has been proposed to operate as a tumor suppressor in some malignancies and a proto-oncogene in others (Shingleton and Hemann, 2015). In addition, we show that *CHD8* haploinsufficiency causes a mild downregulation of Cbl-b transcription — an E3 ligase implicated in degradation of the EGFR — and a marked upregulation of Erk phosphorylation, indicating a repressive action of CHD8 on MAPK activity. The small decrease in Cbl-b mRNA levels (20%) does not scale up with the potent increase in Erk phosphorylation observed in *CHD8*^*+/−*^ suggesting that CHD8 interferes with MAPK signaling in multiple ways. The MAPK pathway operates in early G1 and is essential for (re)-entry into mitosis (Liu et al., 2004). Upregulation of MAPK activity in *CHD8*^*+/−*^ may therefore also drive cell cycle re-entry in a subset of quiescent NPCs, further accelerating G1 in mutant progenitors.

Collectively, our findings provide compelling evidence for dysregulated proliferation of neural progenitors in *CHD8*^*+/−*^ developing brains and strengthens the unexpected connection between autism and cancer (Crawley et al., 2016). Several other autism risk factors are also associated with cancer, including PTEN, and components of the mTOR and Ras-MAPK pathways. PTEN-ASD is estimated to represent up to 2% of all autism cases (Crawley et al., 2016) and is also strongly associated with macrocephaly (Varga et al., 2009). Of note, 27% of people with RASopathies — a group of 5 neurodevelopmental disorders caused by mutations in the Ras-MAPK pathway that all result in Erk gain-of-function — meet the criteria for autism (Adviento et al., 2014). Our work thus identifies MAPK signalling as a potential therapeutic target for the treatment of the CHD8 ASD subtype and suggests that cancer drugs targeting the MAPK pathway may be therapeutically active in idiopathic autism.

## Acknowledgments

We thank Prof. Lauren Pecorino, Dr Yvonne Walsh and Patricia Lopez Garcia for critical reading of the manuscript. We are very grateful to Dr. Andy Bashford (Molecular Devices) for giving us access to the ImageExpress MicroConfocal microscope for some of our imaging experiments. This work was supported by a project grant from the Leverhulme Trust (RPG-2018-265) to MF.

