## Supplemental Material for "Autism-associated CHD8 keeps proliferation of human neural progenitors in check by lengthening the G1 phase of the cell cycle"

### Supplemental data

**Figure S1. Surveyor assay and TIDE results.** (a) Surveyor assay performed on hESCs transfected with pSpCas9-ex3-gRNA and pSpCas9-ex4-gRNA. The cleavage pattern observed for both gRNAs (red asterisks) is consistent with a double strand break at the predicted site. The predicted positions of the DSB sites in these genomic amplicons are depicted below. These cleavage products are not observed in hESCs transfected with GFP. (b) Predicted allele frequency and composition of the E11 hESC clone by TIDE based on sequencing results of a genomic amplicon.

**Figure S2.** (a) Image of a hESC colony transduced with FUCCI. The green and red channels are overlaid on top of a brightfield image. (b,c) Snapshots of FUCCI-transduced *CHD8*$^{+/+}$ and *CHD8*$^{+/-}$ hESCs taken over the course of ~ 20hrs. The white dotted line shows expansion of the stem cell colony over time. Time is indicated in hours and mins. Scale bar: 20 μm.

Figure S1 Coakley-Youngs et al.,

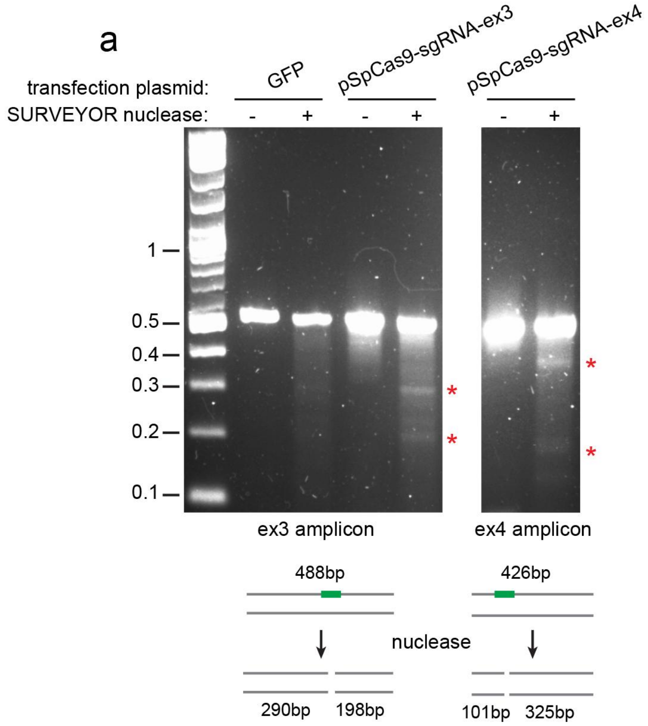


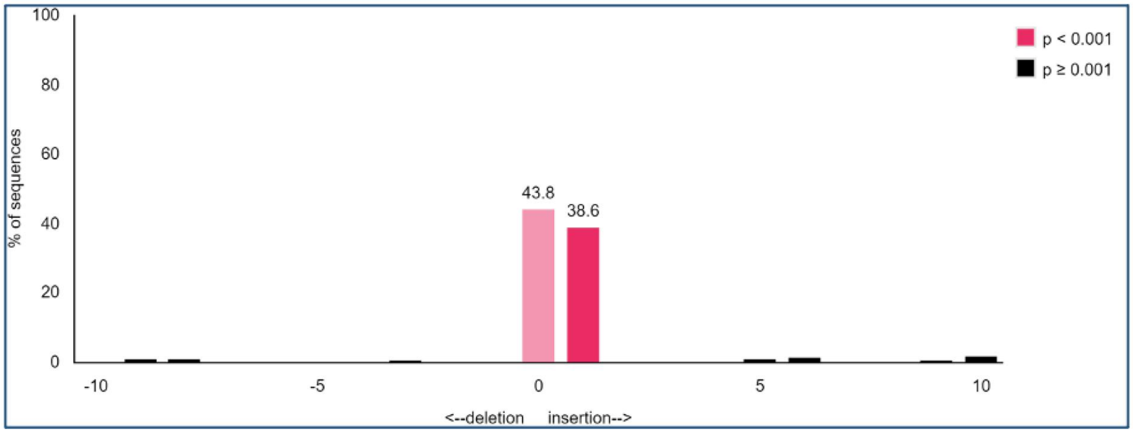

Figure S2 Coakley-Youngs et al.,

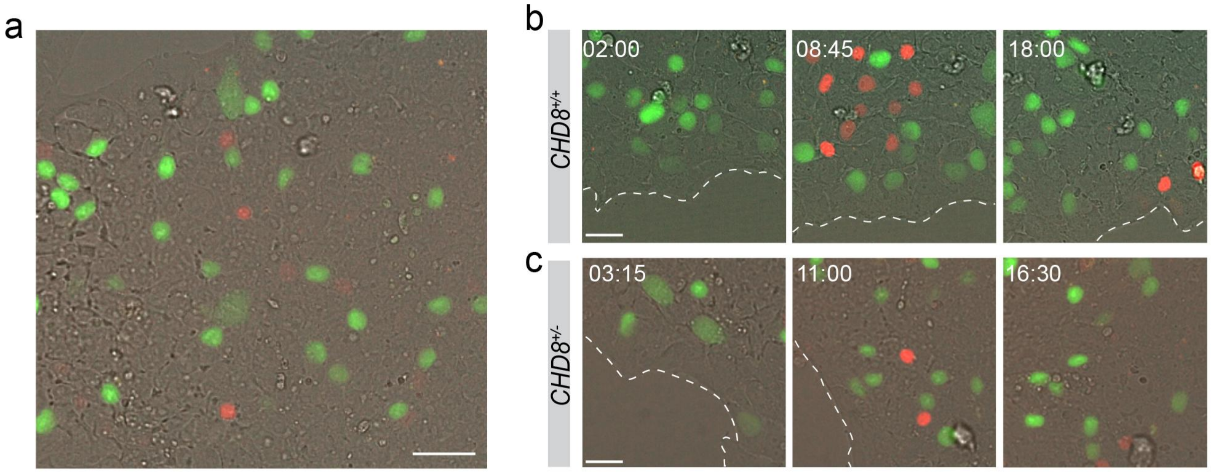

### Matlab script

```
% This script uses a point-and-click strategy to segment and track FUCCI-labeled cells
% in a time series.
% The script displays the read and green FUCCI intensities (C1Int and C2Int) for all
% cells identified in a lineage.
% The scripts saves a spreadsheet containing coordinates of each click and
% corresponding FUCCI intensities in both channels.
% last modified by Marc Fivaz, October 2020.

clear all, close all
```

#### load up stacks

```
C1 = 'R-G07.tif'; % change this to the name of your red stack
InfoImage=imfinfo(C1);
mImage=InfoImage(1).Width
nImage=InfoImage(1).Height
NumberImages=length(InfoImage)

FinalImage=zeros(nImage,mImage,NumberImages,'uint8');
for i=1:NumberImages
  C1s(:,:,i)=imread(C1,'Index',i);
end

C2 = 'G-G07.tif'; %change this to the name of your green stack
InfoImage=imfinfo(C2);
mImage=InfoImage(1).Width;
nImage=InfoImage(1).Height
NumberImages=length(InfoImage)

FinalImage=zeros(nImage,mImage,NumberImages,'uint8');
for i=1:NumberImages
  C2s(:,:,i)=imread(C2,'Index',i);
end
```

#### local background subtraction

```
se = strel('disk', 30);
for k = 1:NumberImages
          C1s_bg(:,:,k) = imopen(C1s(:,:,k), se);
          C1s_corr(:,:,k) = C1s(:,:,k) - C1s_bg(:,:,k);
          C2s_bg(:,:,k) = imopen(C2s(:,:,k), se);
          C2s_corr(:,:,k) = C2s(:,:,k) - C2s_bg(:,:,k);
end
```

#### variables

```
filename = 'G07_s2';
timeInterval = 915.86; % extract from meta data
FrameNumber = NumberImages;
time = [0:15.264:(NumberImages-1)*15.264];
time = time';
```

### binary image from image from each channel

```
for i = 1:NumberImages
        TC1 = adaptthresh(C1s_corr(:,:,i), 0.15); % adaptive thresholding
        TC2 = adaptthresh(C2s_corr(:,:,i), 0.075); % adaptive thresholding
        bwC1(:,:,i) = imbinarize(C1s_corr(:,:,i), TC1);
        bwC2(:,:,i) = imbinarize(C2s_corr(:,:,i), TC2);
        bwC1(:,:,i) = imfill(bwC1(:,:,i), 'holes');
        bwC2(:,:,i) = imfill(bwC2(:,:,i), 'holes');
        bwC1(:,:,i) = bwareaopen(bwC1(:,:,i), 50); % minimum size
        bwC2(:,:,i) = bwareaopen(bwC2(:,:,i), 50); % minimum size
end
```

### sum C1 and C2 binary stacks and visualize segmentation

```
for i = 1:NumberImages
        bw_sum(:,:,i) = bwC1(:,:,i) + bwC2(:,:,i);
        bw_sum_perim(:,:,i) = bwperim(bw_sum(:,:,i));
        ImMerge = imfuse(C1s_corr(:,:,i), C2s_corr(:,:,i), 'ColorChannels', [1 2 0]);
        ImOv(:,:,:,i) = imoverlay(ImMerge, bw_sum_perim(:,:,i), [1 1 1]);
end
```

### interactive segmentation of cells and their progeny

```
 for i = 1:NumberImages
        C1s_corr_adj(:,:,i) = imadjust(C1s_corr(:,:,i), [0.005 0.02], []);
        C2s_corr_adj(:,:,i) = imadjust(C2s_corr(:,:,i), [0 0.07], []);
        ImMerge = imfuse(C1s_corr_adj(:,:,i), C2s_corr_adj(:,:,i), 'ColorChannels', [1 2 0]);
        imshow(ImMerge, []), tqmax;
        C11s = C1s_corr(:,:,i);
        C22s = C2s_corr(:,:,i);
        [col,row] = ginput;

        for k = 1:length(col)
        bw2 = bwselect(bw_sum(:,:,i), col(k), row(k), 8);
        imshow(bw2);
        pause(0.05)
        CC = bwconncomp(bw2, 8);
        TF = isempty(CC.PixelIdxList);
        if TF == 1
```

```
            C1Int(i,k) = mean(mean(C1s_bg(:,:,i)));
            C2Int(i,k) = mean(mean(C2s_bg(:,:,i)));
            elseif TF == 0
            C1Int(i,k) = mean(C11s(CC.PixelIdxList{1,1}));
            C2Int(i,k) = mean(C22s(CC.PixelIdxList{1,1}));
            end
            RInt(i,k) = C1Int(i,k);
            GInt(i,k) = C2Int(i,k);
            end
            allcoord2(:, 1) = col;
            allcoord2(:, 2) = row;
            allcoord2(:, 3) = 1:length(col);
            allcoord2(:, 4) = i;
            allcoord2(:,5) = C1Int(i,:)';
            allcoord2(:,6) = C2Int(i,:)';
            if i == 1
            allcoord = allcoord2;
            else
            allcoord = vertcat(allcoord, allcoord2);
            end
            clear allcoord2;
            clear C1Int;
            clear C2Int;
 end
```

#### Plot intensity traces

```
figure, hold on
plot(time, GInt(:,1), 'g');
plot(time, RInt(:,1), 'r');
hold off
xlabel('time min');
ylabel('Int');
legend('S/G2/M', 'G1');
title('cell 1')

figure, hold on
plot(time, GInt(:,2), 'g');
plot(time, RInt(:,2), 'r');
hold off
xlabel('time min');
ylabel('Int');
legend('S/G2/M', 'G1');
title('cell 2');
```

```
% repeat if lineages includes more than two daughter cells
```

### create and export dataset

```
T = table(allcoord (:, 1), allcoord (:, 2), allcoord (:, 3), allcoord(:, 4), allcoord(:, 5), allcoord(:, 6));
T.Properties.VariableNames = {'column' 'row' 'cellnumber' 'frame' 'C1Int' 'C2Int'};
fname = strcat(filename, '_CoordInt.xlsx');
writetable(T, fname);
```
